# Uncoupling microtubule lifetime, stability and post-translational modifications

**DOI:** 10.64898/2026.09.10.750621

**Authors:** Tanguy Chocat, Benoit Vianay, Flora Silberzan, Jérémie Gaillard, Louise Bonnemay, Alexandre Schaeffer, Laurent Blanchoin, Manuel Théry

## Abstract

Microtubules (MTs) undergo continuous cycles of growth and disassembly. Because the transitions between these states are stochastic, MT age varies widely within a population. As MTs age, they are thought to accumulate post-translational modifications (PTMs) that, directly or indirectly, enhance their stability and thereby extend their lifetime. The rare MTs that withstand prolonged exposure to destabilizing drugs such as nocodazole (NZ) are indeed enriched in PTMs; yet the relationships between MT age, PTMs and stability remain unclear. Using microinjection of labelled tubulin, we measured microtubule network turnover in immortalized mouse embryonic fibroblasts. Half of the network was renewed within 3 minutes and 80% within 10 minutes, while approximately 5% of microtubules persisted for more than 20 minutes. These dynamics were comparable in quiescent and senescent cells, although the fraction of slowly renewing or non-renewing microtubules rose to 20% in senescent cells. Unexpectedly, neither the amount of PTMs (acetylation and detyrosination) nor resistance to NZ increased with MT age, and resistance to NZ was independent of these PTMs. Degrees of acetylation and detyrosination should therefore not be taken as readouts of MT age or stability. Because these PTMs do not accumulate on MTs over time, the chemical modification of polymerized tubulin is likely more reversible and dynamic than previously assumed.

## Introduction

Microtubules (MTs) grow and shrink continuously, at rates that vary between cell types (Wadsworth and McGrail, 1990). In fibroblasts, for example, they grow at roughly 5-10 µm/ min and can stochastically stop growing and depolymerize at 10-20 µm/min, so that the whole network is renewed within 5 to 20 minutes (Saxton et al., 1984; Schulze and Kirschner, 1986; Cassimeris et al., 1988). This renewal rate varies with cell-cycle progression (Belmont et al., 1990; Piehl et al., 2004; Zhai et al., 1995), quiescence (Laporte et al., 2015) and differentiation (Wehland and Weber, 1987; Lacroix et al., 2014; Fraser et al., 2025). Such variations likely reflect changes in the concentrations of the many microtubule-associated proteins (MAPs) that govern the addition, maintenance and removal of tubulin at MT ends (Akhmanova and Steinmetz, 2015).

Notably, some MTs within the network are less dynamic than others, pausing and depolymerizing less frequently (Cassimeris et al., 1988). In BSC1 fibroblasts, a further subset is not dynamic at all, retaining their tubulin for more than an hour (Schulze and Kirschner, 1987). These so-called “stable MTs” have been proposed to play a key role in specifying the polarity axis of migrating or differentiating cells (Etienne-Manneville, 2013; Baas et al., 2016; Muroyama and Lechler, 2017; Scarborough et al., 2026) which makes understanding the origin and function of these subsets particularly important (Jansen et al., 2023).

MT stability is characterized using various terms and readouts including MT lifespan, resistance to perturbations, and the presence of post-translational modifications (PTMs) on tubulin (Janke and Magiera, 2020; Roll-Mecak, 2020; Verhey and Gaertig, 2007). These three parameters, however, capture quite distinct properties that should be kept separate, and much of the confusion in the field stems from using the single term “stable” for all of them. “Stable” means able to resist perturbations such as tubulin dilution or mechanical stress, but does not imply that, in the absence of perturbation, such microtubules would live longer (much as a helmet improves a cyclist’s odds in a crash without extending their life expectancy otherwise). “Long-lived” means that a microtubule’s intrinsic life-time is increased but does not imply resistance to perturbation (much as a healthy lifestyle offers no protection in a bicycle accident). Distinct assays have been designed to probe each of these features.

Resistance to perturbation, that is true stability, is typically assayed by exposure to cold or to tubulin-sequestering compounds such as nocodazole (NZ) (Bré et al., 1987). The few MTs that survived prolonged NZ treatment (an hour or more) were shown to be highly detyrosinated and acetylated (Bré et al., 1987; Wehland and Weber, 1987; Kreis, 1987; Piperno et al., 1987). MT lifetime in contrast, is measured by microinjecting labelled tubulin and following its incorporation into newly assembled microtubules (Schulze and Kirschner, 1986), which allows new MTs to be distinguished from older ones that have not incorporated the injected tubulin (Schulze and Kirschner, 1987). Intriguingly and easily confused with NZ-resistance experiments, the few MTs that had not renewed their tubulin after more than an hour were likewise highly detyrosinated and acetylated (Webster et al., 1987; Schulze et al., 1987b; Webster and Borisy, 1989). The three readouts – age, resistance and modifications – were thus merged and treated as as interchangeable measurements of MT “stability”. This equivalence was reinforced by the functional association of the three in migrating cells during wound healing, where MTs were described as being captured at the cell front, which increased their lifetime and their potential for detyrosination (Gundersen and Bulinski, 1988). Because tubulin modification was considered as slow process taking 20-30 min to appear on MTs regrowing after NZ washout (Bré et al., 1987; Gundersen et al., 1987; Khawaja et al., 1988), and because detyrosination and acetylation favour the recruitment of kinesins (Liao and Gundersen, 1998; Reed et al., 2006), that could transport factors to the cell front, a positive feedback loop was proposed to polarize migrating cells by generating a sub-population of “stable MTs” that are simultaneously long-lived, modified and NZ-resistant (Li and Gundersen, 2008).

Yet, the equivalence between these three features should not be assumed to be systematic, for several reasons. First, chemical diversity: different tubulin PTMs can mark distinct subsets of MTs within the same cell (Bulinski et al., 1988), so no single PTM can, on its own, be part of a common stabilization process and each modified subset may have its own dynamics and stability properties. Second, molecular function: despite the correlations observed in cells, it soon became clear that detyrosination or acetylation do not, per se, prolong MT lifetime (Webster et al., 1990; Schulze et al., 1987a) or improve NZ resistance (Khawaja et al., 1988); indeed, in vitro reconstitution later showed that neither modification affects MT dynamic parameters (Portran et al., 2017; Chen et al., 2021). Third, confounding secondary effects: tubulin PTMs can promote the recruitment of MAPs and motors and thereby influence MT stability indirectly (Peris et al., 2006; Chen et al., 2021), but the possible combinations and competitions among tubulin isotypes, PTMs, MAPs and motors are so varied that they can produce almost any consequences for MT dynamics and stability (Monroy et al., 2020; Roll-Mecak, 2020). Fourth, and central to our study, the time scale: the early use of one-hour window to identify non-renewing or NZ-resistant MTs is very long relative to the ∼10 min needed to renew most of the network, and the functional impact of tubulin PTMs on the subset of dynamic MTs has not been tested. Acetylation, for instance, can be detected in cells displaying rapid turnover MTs (Schulze et al., 1987a), and on the short-lived microtubules of the mitotic spindle (Barisic et al., 2015; Velasquez-carvajal et al., 2024), showing that tubulin modification can be rapid and is not specific to long-lived microtubules. The widely accepted correlation between MT age, modification and stability may therefore be more complex than initially described, particularly for the large, rapidly turning-over fraction of the network.

Here we revisit these relationships by reviving the early tubulin-injection method of the Kirschner laboratory to distinguish renewed and non-renewed microtubules (Schulze and Kirschner, 1987), allowing us to measure MT age and relate it, through fine temporal dissection, to the extend of tubulin modifications and to resistance to nocodazole.

## Results and discussion

### Measurement of MT age and network turnover

We worked with immortalized mouse embryonic fibroblasts (MEF) (Virtakoivu et al., 2015). Microinjection of labelled tubulin reveals the new microtubules that assemble between injection and cell fixation (Saxton et al., 1984; Schulze and Kirschner, 1986; Cassimeris et al., 1988; Gazzola et al., 2023). However, these new microtubules are also built from endogenous tubulin, which is likewise present in older MTs that were not renewed during this interval (Fig. 1A). The old, pre-existing MTs therefore cannot be observed in a dedicated fluorescent channel and are often masked by the new microtubules in the dense regions near the cell center (Fig. 1B). To overcome this limitation, the Kirschner laboratory devised a protocol to shield the new microtubules after fixation and stain the old ones specifically (Schulze and Kirschner, 1987). The injected biotinylated tubulin was labelled with goat anti-biotin antibodies, which were then bound by bovine anti-goat antibodies, themselves bound by goat anti-bovine antibodies. Iterating these steps coated the biotin-containing microtubule with several dense antibody layers that act as a shield, preventing other antibodies from accessing their lattice (Fig. 1C). Any subsequent immunostaining was thus restricted to the non-renewing microtubules, which had not incorporated the injected tubulin and could therefore be identified specifically (Fig. 1C, D).

**Figure 1.**
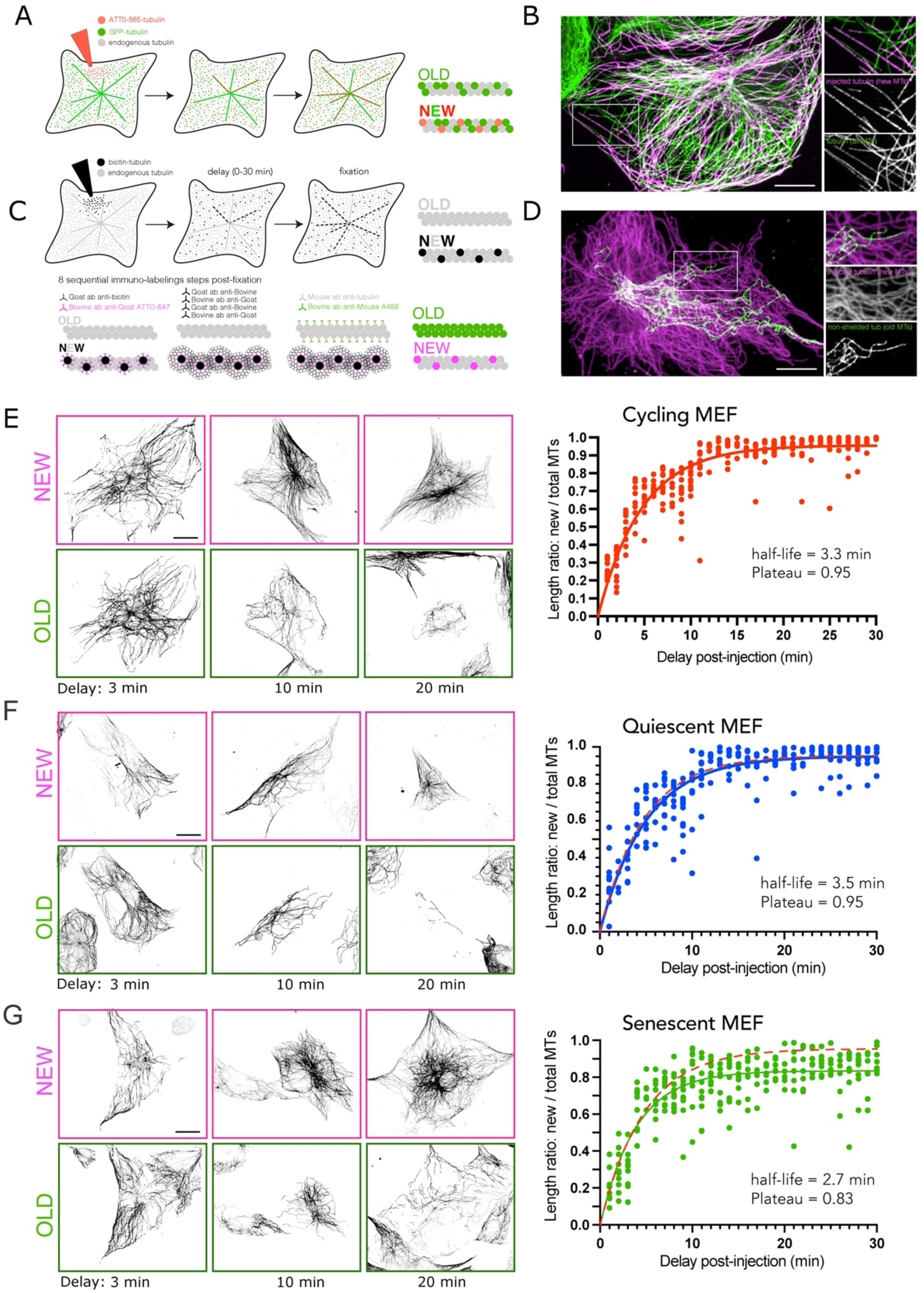
Turnover of the MT network. (A) Schematic of the tubulin injection protocol revealing newly formed MTs. (B) Example showing the MTs composed of fluorescently-labelled injected tubulin (magenta) and the entire pool of MTs (green). Scale bar corresponds to 10 µm. (C) Schematic of the tubulin injection protocol allowing the distinction between newly formed (« NEW ») and pre-existing (« OLD ») MTs. The shielding of « new » MTs containing injected tubulin with several layers of antibodies allows the selective immuno-staining of « old » MTs, which existed before the injection and therefore do not contain the injected tubulin (D) Example showing the MTs composed of biotinylated, injected and shielded tubulin (magenta) and the pre-existing and non-shielded MTs (green). Scale bar corresponds to 10 µm. (E) Images show examples of « new » MTs, assembled during the delay after tubulin injection, and « old » MTs, pre-existing to the injection and maintained during the delay, for 3, 10 and 20 minutes long delays in cycling MEF. In each condition, « new » MTs are younger than the delay and « old » MTs are older than the delay. Scale bar represents 10 µm. The graph represents the quantification of the ratio of the lengths between the « new » MTs and the entire network made of « new » and « old » MTs for various delays between the injection and fixation of cells. The curve thus represents the time-scale of replacement of « old », pre-existing MTs, by « new » MTs. Values were fitted with a one-phase decay exponential. (F) Same as (E) in quiescent cells, cultured in low serum (0.5%) for two days. (G) Same as (E) In senescent cells, treated for five days with 12 µM of palbociclib.

Throughout this report, we refer to “new” MTs as those assembled between injection and fixation (magenta in Fig. 1D), and to the “old” MTs as those present at the time of injection that had not disappeared before fixation (in green in Fig. 1D). This method does not reveal the exact age of MTs but categorizes it: “new” MTs are younger than the delay between injection and fixation, and “old” MTs are older than it. By varying this delay between 0 and 30 minutes, we could follow the evolution of the relative proportion of renewed (“new”) and non-renewed (“old”) MTs over time (Figure 1E, Fig. S1A). Plotting the proportion of new MTs relative to the entire pool (new plus old) and fitting it to a one-phase-decay process, we found that half of the MTs were renewed in 3 minutes and about 80% within 10 minutes (Fig. 1E). Roughly 5% of the network remained non-renewed after 20 minutes (Fig. S1B). As previously described, these long-lived MTs appeared curlier than young MTs, but they were not all associated with the nucleus (Fig. S1B).

### MT network turnover in quiescent and senescent cells

Earlier work on MDCK cells showed that MT network turnover is dramatically reduced during the epithelial morphogenesis, as cells form intercellular contacts and polarize (Bré et al., 1987; Bre et al., 1990). We therefore investigated how exit from the cell cycle, a key step in cell differentiation, influences MT network turnover, testing two forms of exit: the reversible shift to quiescence and the irreversible progression to senescence. Quiescence was induced by lowering the amount of fetal calf serum in the culture medium from 10 to 0.5%. Two days later, almost all cells were quiescent, as shown by the presence of primary cilia (Fig. S2A). Surprisingly, injection of biotin-labelled tubulin revealed that MT turnover in these quiescent cells was almost unaffected (Fig. 1F). Notably, cilia were no longer visible in the non-renewing MT population after 20 minutes, suggesting that the MTs of primary cilia had been renewed (Fig. S2B).

Senescence was induced in MEF by adding 12 µM palbociclib, a selective inhibitor of CDK4 and 6 (Michaud et al., 2010), to the culture medium. Five days later, all cells were positive to β-galactosidase, a well-characterized marker of complete senescence (Dimri et al., 1995) (Fig. S2C). In these cells, the tubulin renewal rate was similar to that of proliferating cells (half of MTs renewed within 3 minutes), but the proportion of non-renewed microtubules after 20 minutes rose from 5 to 20% (Fig. 1G and S2D). These data indicate that exit from the cell cycle does not entail a slowdown of MT dynamics, but that senescence immobilizes a larger subset of non-renewing MTs.

### Tubulin modifications in old and new MTs

We next investigated the extent of tubulin modifications in pre-existing (“old”) and renewed (“new”) MTs. Because, the lattice shielding of the lattice used to distinguish old from new MTs blocks antibodies to new-MT lattice, immunostaining of tubulin PTMs had to be performed before the shielding step (Fig. 2A). We could thus visualize tubulin detyrosination and acetylation, together with the old and new MTs subsets, in four dedicated fluorescent channels (Fig. 2B). However, these networks overlapped at the resolution of fluorescence microscopy, making it difficult to assign each modification specifically to old or new MTs. We therefore developed an image analysis pipeline to improve the accuracy of our PTM estimates for each subset.

**Figure 2.**
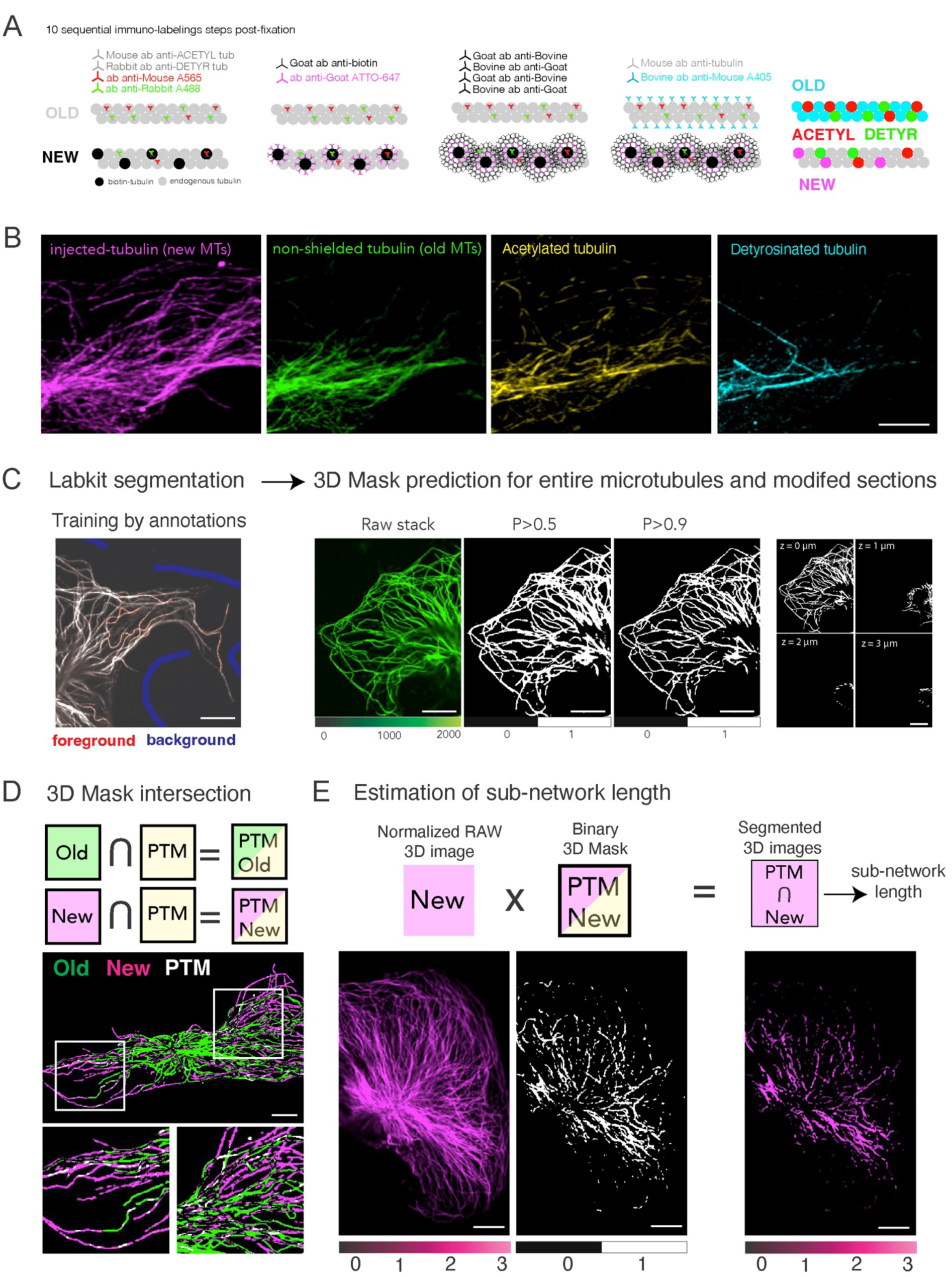
Segmentation of MTs and quantification of tubulin post-translational modifications (PTMs). (A) Schematic of the immuno-labelling protocol allowing the detection of tubulin acetylation and detyrosination as well as the age of MTs with the tubulin injection and shielding strategy. (B) Examples of images showing the combination of stainings for old and new MTs, acetylated and detyrosinated tubulins. Scale bar represents 10 µm. (C) Illustration of the segmentation protocol. The training step, based manual annotations of MTs, allowed Labkit to automatically predicts whether pixels belong to MTs or to the background with a given probability. Varying this value, the MT network could be segmented in 3D with various degree of confidence. The segmentation led to a 3D binary mask. (D) Calculation of the intersection of masks produced from stainings of PTM and from stainings of old or new MTs led to 3D mask of pixels specific to old-and-modified (acetylated or detyrosinated) or new-and-modified sections of MTs. (E) Calculation of the length of modified networks. 3D Images of new or old MTs were normalized by the intensity of single MTs so that the intensity of each pixel correspond to the number of MTs. The multiplication of these stacks (containing the number of MTs per pixel) with the old/new-and-modified masks (specifying which pixels correspond to new or old, modified or not, part of the network) led to the estimation of the total length of the corresponding sub-networks.

We used LabKit to classify pixels and segment MTs (Arzt et al., 2022). The algorithm was trained on our images by annotating linear regions as either MT or background. By adjusting the probability threshold for a pixel to belong to a MT, we could obtain more or less restrictive segmentations (Fig. 2C). A low probability threshold (P>0.5) gave continuous segmentation that closely followed the linear shape of MTs, whereas a higher probability threshold (P>0.9) produced graphically less satisfying shapes but ensured the specificity of pixel classification, which we preferred for quantification (Fig. 2C). Each plane of Z stacks was segmented to generate 3D binary masks corresponding to the network’s 3D footprint. We thereby produced masks for old and new MTs as well as for acetylated and detyrosinated MT segments. To estimate the degree of modifications on old and new MTs, we computed the intersections of the age and modifications masks (Fig. 2D). We then needed to count the number of modified MTs in each pixel of these masks. To do so, the pixels intensities of the raw old and new network 3D stacks were normalized to the intensity of single MTs, so as to reflect the number of MTs per pixel. Multiplying the intersection mask by this per pixel MT-number stack yielded a stack in which each pixel’s intensity corresponded to the number of modified MTs (Fig. 2E). Summing the intensity of all pixels then gave an estimate of the total modified length within old and new networks.

Given the optical resolution (300nm) and the pixel size (150nm) relative to the MT diameter (25 nm), however, old and new MTs could still fall within the same pixel, making it difficult to assign a PTM signal to one network or the other. We therefore excluded all regions of overlap between the old and new masks to ensure that the PTMs we quantified lay exclusively on one subset (Fig. S3A). We could verify visually that the segmented modified segments corresponded unambiguously to either old or new MTs, with no overlap (Fig. S3A).

We could thus reliably compare the proportions of modification on new and old MTs at various delays after injection (Figure 3A). Surprisingly, the amount of acetylation did not increase with MT age and even appeared independent of it (Fig. 3B). Four minutes after injection, about 10 % of the length of young MTs (ie younger than 4 minutes) was acetylated (Fig. 3B, C). We measured similar proportions on old MTs, (ie on MT older than 5 minutes), and these proportions held at longer delays as well: twenty-five minutes after injection, the few MTs older than 25 minutes showed no greater acetylation (Fig. 3B, D). Detyrosination was more difficult to quantify across many cells (Fig. 3E), because only about 60% of MEF had detyrosinated MTs (see Fig. 5D for a wide field of view). Considering that in general many cells suffered from the injection and had to be discarded, only 20% the cells we injected cells could be analyzed. Nonetheless, it was clear that detyrosination did not increase with MT age (Fig. 3F) and remain restricted to about 5% of MT length. As with acetylation, some young microtubules were detyrosinated (Fig. 3G) and some old ones were not (Fig. 3H). Excluding PTM quantification on the old/new overlap region removed a substantial fraction of modified segments, but even when these were included no correlation between PTM level and age was observed (Fig. S3B-E).

**Figure 3.**
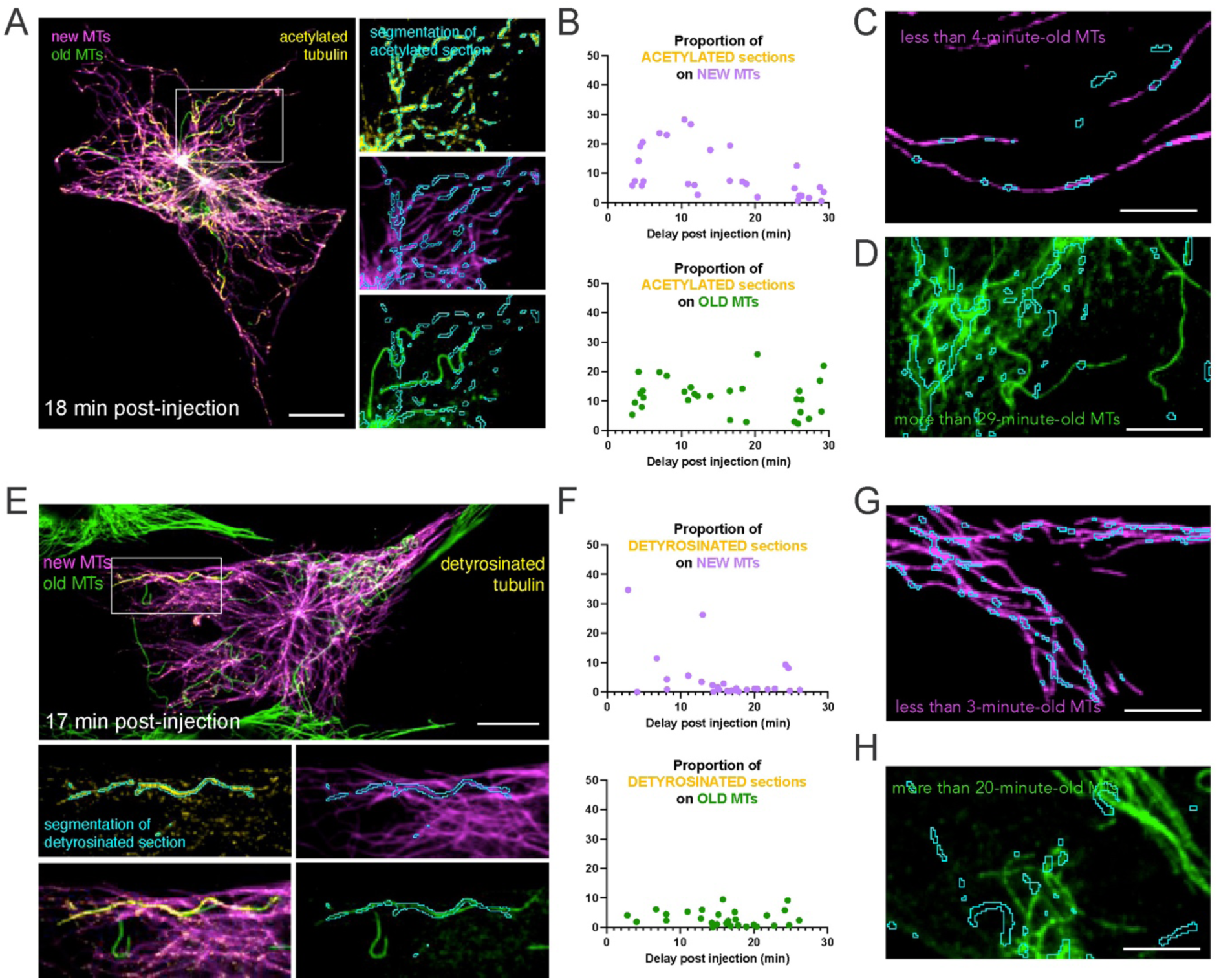
Relationship between MT age and tubulin post-translational modifications (PTMs). (A) Example of the segmentation of the pattern of acetylated tubulin in a cell fixed 18 minutes after tubulin injection. The top magnified inset shows the regions of interest (blue contours) around acetylated sections (yellow). These regions have been reported on channels corresponding to injected tubulins (magenta), ie MTs younger than 18 minutes, and shielded tubulins (green), ie MTs older than 18 minutes. Scale bar corresponds to 10 µm. (B) Graphs show the length of the « new » and « old » networks that were acetylated relative to the entire « new » and « old » network, depending on the delay after injection, ie the age limit between these two populations. (C) An example showing the presence of acetylated sections on young MTs, which are less than 4-minute-old. (D) Another example showing the opposite, ie their absence on some old MTs, which are more than 29-minute-old. Scale bar corresponds to 5 µm. (E) Example of the segmentation of the pattern of detyrosinated tubulin in a cell fixed 17 minutes after tubulin injection. The top magnified inset shows the regions of interest (blue contours) around detyrosinated sections (yellow). These regions have been reported on channels corresponding to injected tubulins (magenta), ie MTs younger than 17 minutes, and shielded tubulins (green), ie MTs older than 17 minutes. Scale bar corresponds to 5 µm. (F) Graphs show the length of the « new » and « old » networks that were detyrosinated, depending on the delay after injection, ie the age limit between these two populations. (G) An example showing the presence of detyrosinated sections on young MTs, which are less than 4-minute-old. (H) Another showing the opposite, ie their absence on some old MTs, which are more than 20-minute-old. Scale bar corresponds to 5 µm.

These data run counter to several widely held hypotheses about the action of modifying enzymes. Our conclusions cannot be generalized, as they come from a single cell type, but they provide counter-examples sufficient to challenge several earlier conclusions. First, tubulin modification within a microtubule can be fast: we detected acetylation and detyrosination on microtubules only a few minutes old. Second, these modifications do not accumulate with age. One possible mechanism could be a process that prevents partially modified MTs from undergoing further modification. Alternatively, the modifications could be reversible, i.e. constantly added to and removed from the tubulin in the lattice, or the modified tubulins could constantly be removed and replaced by non-modified tubulins. In the later cases, the extent of modifications on a MT would reflect not its age but the local balance between modifying and demodifying processes in its immediate environment.

### Stability of old and new MTs

To characterize MT stability, we tested their resistance to free tubulins sequestration upon addition of 10 µM of nocodazole (Figure 4A). Cells were fixed after various delays and the length of the remaining network was estimated with our image-analysis pipeline. As expected, the network disappeared progressively. Fitting the data as a one-phase-decay process, we obtained exactly the same parameters as for network renewal: half of the population disappeared within 3 minutes and 80% within 10 minutes (Figure 4A). NZ addition therefore revealed the unperturbed dynamics of MT disassembly and merely prevented MT reassembly. The kinetics of MTs disassembly under NZ can thus serve as a reliable readout of network turnover.

**Figure 4.**
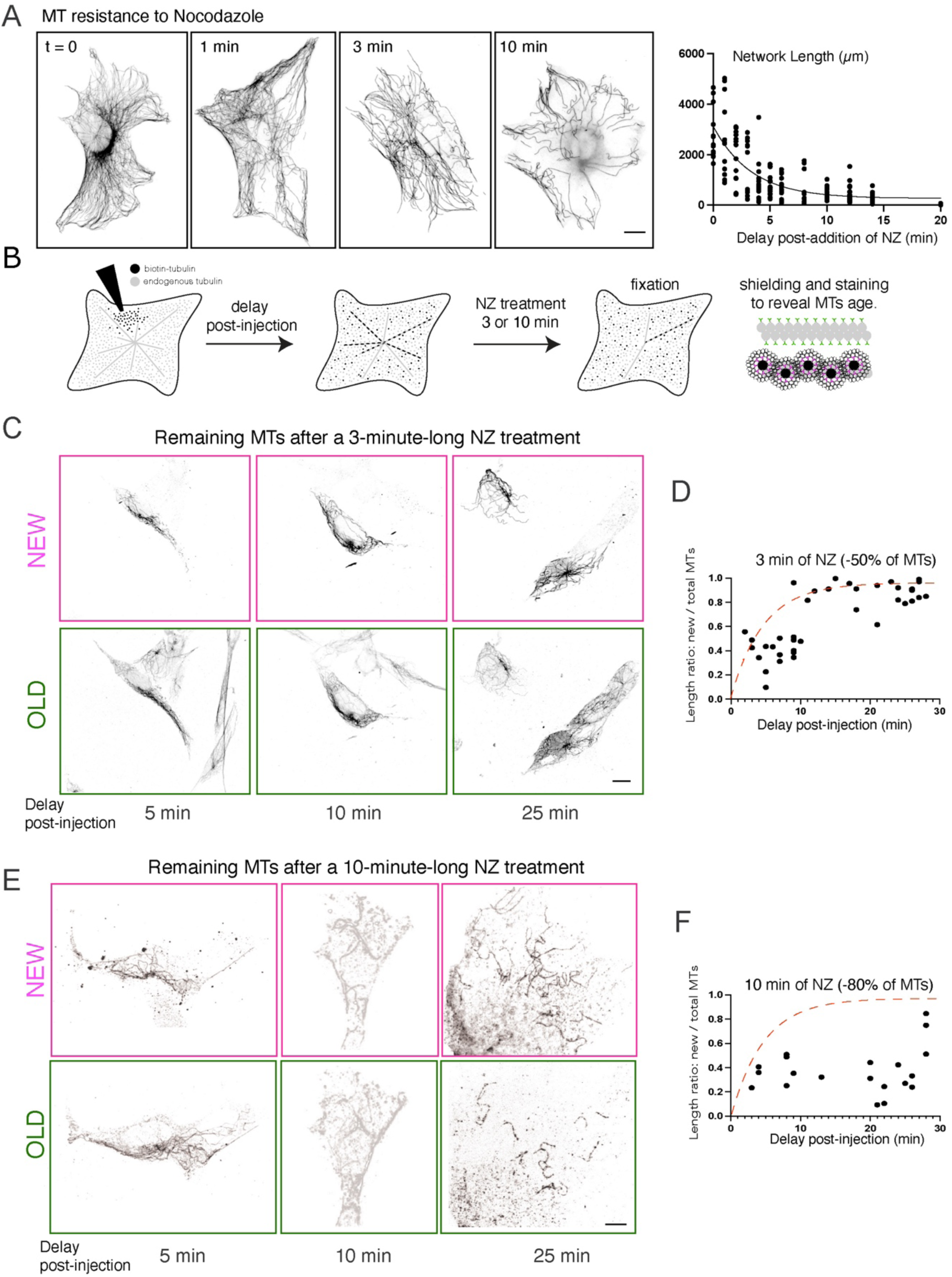
Relationship between MT age and stability. (A) Assessment of MTs stability by their resistance to 10 µM Nocodazole (NZ). Images show MT immuno-staining following cell exposure to 10 µM NZ for various delays. Graph shows the quantification of MT network length with respect to the duration of NZ exposure. Values were fitted with a one-phase decay exponential. (B) Schematic of protocol combining tubulin injection, to measure MT age, and exposure to NZ, to assess MT stability. Depending on the delay after injection, the age limit between old and new MTs, as well as their relative proportions were varied. These populations of old and new MTs were then exposed to NZ. Two durations of NZ treatment were used, 3 and 10 minutes, in order to disassemble 50% and 80% of the network. Fixation was followed by the shielding protocol (see Figure 1C) to distinguish old and new MTs. (C) Images show the remaining old and new MTs following 3-minute exposure to 10 µM NZ. The delay post-injection indicates the age limit between old and new MTs. Scale bar corresponds to 10 µm. (D) Graph shows the remaining proportion of new MTs relative to the total network after 3 minutes exposure to 10µM NZ. The dotted red line is the fit of the values in non-treated cells (shown in Figure 1E). (E, F) Same as (C, D) after 10 minutes exposure to 10 µM NZ.

We next applied 3– and 10-minute exposure to 10 µM NZ to disassemble 50% or 80% of the network, and measure the proportion of new and old MTs among the resistant subsets (Fig. 4B). NZ was added at various delays after tubulin injection, to determine how the initial ratios changed upon disassembly (Fig. 4C). The evolution of these ratios with injection time was similar before and the loss of 50% of the network, indicating that both old and new MTs disassembled in the same proportions (Fig. 4D). After the loss of 80% of the network, the ratios shifted toward higher proportions of old MTs (Fig. 4E, F). However, they did not approach zero, showing that resistant MTs were not exclusively old and non-renewing.

These data indicate that the disassembly is stochastic and not specific to the oldest part of the dynamic MT pool. Moreover, NZ resistance does not identify old non-renewing MTs alone: The observation that some young, few-minutes old MTs could resist NZ (Fig. 4C and E, left columns), suggests that protecting MAPs can also be loaded soon after MT assembly.

### Tubulin modifications of stable MTs

Having uncoupled MT age from both tubulin-modification content and NZ resistance, we asked whether these last two parameters were related. The amount of tubulin modification, acetylation or detyrosination, is often used as a proxy for MT stability (Janke and Magiera, 2020; Roll-Mecak, 2020; Verhey and Gaertig, 2007), but this rationale rests on early studies that considered only long NZ treatments and the few MTs that resisting them (Webster et al., 1987; Schulze et al., 1987b; Webster and Borisy, 1989). Here we tested the capacity of PTMs to slow disassembly and confer NZ resistance by exposing MEFs to 10 µM NZ for 3, 10 and 30-minutes, and revealing the PTM content of the remaining MTs by immunostainings (Fig. 5A).

**Figure 5.**
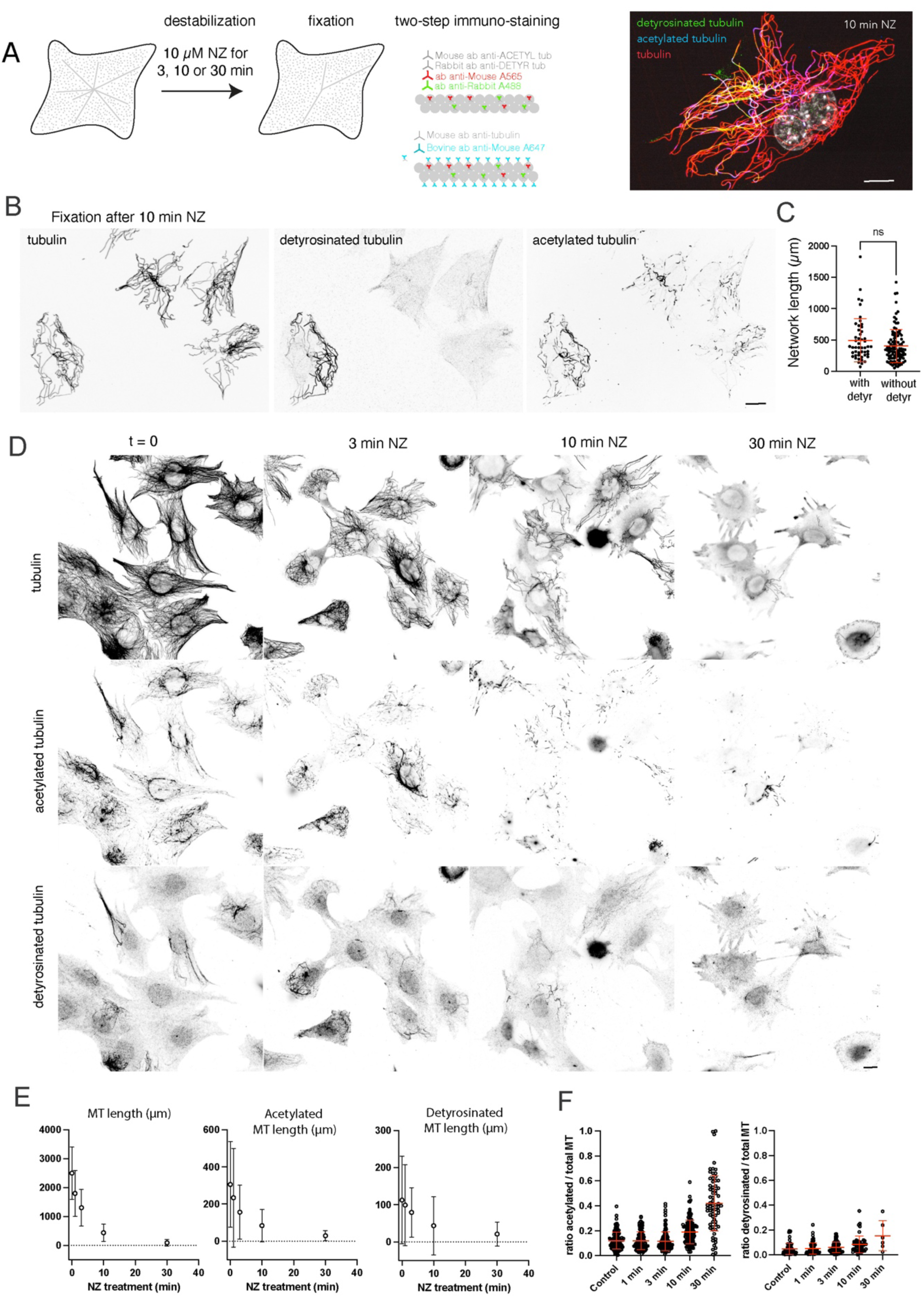
Relationship between MT stability and tubulin post-translational modifications (PTMs). (A) Schematic of protocol to measure MT stability, ie their resistance to NZ, and the localization and density of PTMs. Cells were exposed to 10 µM NZ for 3 to 30 minutes and then fixed and immuno-labelled to detect acetylated and detyrosinated tubulins. Image shows an example of PTM stainings (detyrosinated on green, acetylated in blue) after a 10-minute-long exposure to NZ. Scale bar corresponds to 10 µm. (B) Images show the localization and density PTMs in cells after a 10-minute-long exposure to NZ. In this example, only one out of the four cells had detyrosinated MTs. The resisting network in this cell does not appear longer than in the others. Scale bar corresponds to 10 µm. (C) Graph reports the length of the MT network after 10 min exposure to NZ in cells displaying detyrosinated MTs or not. (D) Images show the localization and density PTMs in cells after various durations of exposure to NZ. Scale bar corresponds to 10 µm. (E) Graphs report the evolution of the length of the entire MT network, the acetylated network and the detyrosinated network with the duration of NZ treatment. (F) Graphs report the evolution of the ratio of the lengths of the acetylated and the detyrosinated network relative to the entire network with the duration of NZ treatment.

As noted above, numerous MT segments were acetylated in all cells but only about 60% of cells (n= 126) contained detyrosinated MTs (Fig. 5B), an opportunity to assess the impact of this modification on NZ resistance. First, the proportion of cells with detyrosinated MTs fell with treatment duration (58% (n=130) after 3 minutes, 33% (n=150) after 10 minutes and 7% (n=68) after 30 minutes), a first indication that MTs in these cells resisted no better than in others. Second, among cells with detyrosinated MTs, the network remaining after 10 minutes of exposure to NZ was not longer than in cells lacking this modification (Fig. 5C). Together, these observations strongly suggest that detyrosination does not improve MT resistance to NZ.

We used our image analysis pipeline to segment the acetylated network, the detyrosinated network and the entire network after each treatment (Fig. 5D). If acetylation or detyrosination protect MTs from NZ-induced disassembly, the length of these modified networks should have remained constant while the rest of the network disappeared. Strikingly, both modified networks also disassembled rapidly (Fig. 5E). To gauge the relative resistance of modified and non-modified portions of the network, we calculated the proportion of the modified length relative to the entire network in individual cells (Fig. 5F). After 3 minutes of exposure to NZ, when half of the initial network had disassembled, the acetylated and detyrosinated proportions were unchanged relative to the starting point, showing that most of the network was not stabilized by these modifications. After 10 minutes, when only 20% of the network remained, the mean acetylated proportion rose slightly but significantly from 12% to 19%, and the detyrosinated proportion from 5% to 8%. These values rose further to 42% and 15% respectively, after 30 minutes when only few MT segments remained (Fig. 5D, F). Even so, these modified, resistant MTs represented only 10% and 20% of the initial acetylated and detyrosinated pools, showing that modification per se was not a cause of, and not even a good indicator of NZ resistance. Interestingly, after 30 minutes the acetylation pattern showed longer, denser stretches than after 3 or 10 minutes, suggesting these resistant MTs were not pre-existing but were instead modified during the treatment, once less MTs were available as enzyme substrate. The increased density of modification could thus be a consequence rather than the cause, of resistance. Furthermore, although the most resistant MTs tended to carry more modifications, the resistant segments were far from fully modified. Resistance to disassembly was therefore not strictly coupled to tubulin modification, which should not be taken as reliable readout of MT stability.

Altogether these data show that in MEF cells MT ageing is accompanied neither by greater tubulin modification nor by greater resistance to monomer sequestering. Furthermore, acetylated and detyrosinated MT segments were not more resistant to monomer sequestration than unmodified segments. Subcellular PTM pattern therefore reflect neither MT age nor stability; more likely, they carry spatial information arising from the local recruitment and competition of modifying and demodifying enzymes. Further investigation into the mechanisms controlling the localization of tubulin-modifying enzymes will be needed to explain the origin and function of these MT subsets.

## Material and Methods

### Cell culture

Mouse embryonic fibroblasts (MEF) (Virtakoivu et al., 2015) were grown at 37°C and 5% CO₂ in DMEM/F-12 (Gibco, 31331028) supplemented with 10% fetal bovine serum (Life Technologies, 10270106) and 1% antibiotic-antimycotic solution (Gibco,15240062). At each passage, cells were detached with TrypLE (Gibco, 12605036). Quiescence was induced by culturing cells in the presence of 0.5% instead of 10% fetal bovine serum for 48 hours. Senescence was induced by adding 12 μM of palbociclib (Sigma, PZ0383) to the normal culture medium for 5 days. β-galactosidase assays were performed with a senescence detection kit (ab65351).

### Tubulin purification

Biotinylated tubulin was prepared as previously described method (Hyman et al., 1991). Microtubules were polymerized from tubulin at 37◦C for 30 min and NHS-Biotin is added for 20 min (2 mM final concentration, ThermoFisher, 21338). The labeling reaction was stopped with potassium glutamate (100 mM final concentration), and the microtubules were sedimented through cushions of BRB80 supplemented with 60% glycerol. The microtubules were resuspended in cold BRB80, then subjected to two further cycles of depolymerization and polymerization before use. The final pellet was resuspended in microinjection buffer (50 mM potassium glutamate, 1 mM MgCl2, pH 6.8), aliquoted into 500 µl plastic tubes, flash frozen in liquid nitrogen and store at –80°C.

### Microinjection

MEFs were injected as previously described (Gazzola et al., 2023). Glass microneedles were pulled from clark borosilicate thin-wall capillaries (30-0050 Harvard Apparatus) using a vertical pipet puller (PN-3, Narishige). Microneedles were manually controlled with an InjectMan 4 micromanipulator (Eppendorf). Microinjections of MEF cells were performed on an inverted microscope (Nikon Ti2 Eclipse) equipped with a Prime BSI Express CMOS camera (Photometrics) and using a Nikon CFI Plan Fluor 40x/0.75 NA dry objective. Compensation pressures were applied with a FemtoJet 4i (Eppendorf) pump. The medium was maintained at 37°C during the whole experiment using a H-301 heating chamber (Okolab). The exact position and time of injection were recorded for each cell. Cells were then fixed or treated with 10 µM nocodazole (sigma, M1404) for 1, 3, 10 or 30 minutes at 37°C before fixation.

### Cell fixation

Injected cells (Figure 1 to 4) were permeabilized for 15 s at room temperature RT in cytoskeleton buffer (10 mM MES, 138 mM KCl, 3 mM MgCl, 2 mM EGTA) supplemented with 10% glycerol and 0.5% Triton-X100 and then fixed for 10 min at RT in cytoskeleton buffer supplemented with 10% glycerol and 0.1% Triton-X100, 0.05% glutaraldehyde, and 4% of paraformaldehyde. Nocodazole-treated cells (Figure 5) were fixed with 0.05% glutaraldehyde, and 4% of paraformaldehyde in cytoskeleton buffer supplemented with 10% sucrose and 0.05% Tween 20 for 10 minutes. In all conditions, aldehyde functions were then reduced using a NaBH4 solution (1mg/ml in PBS) for 10 minutes at RT. The samples were then washed 3 times with PBS-Tween 20 (1379, Sigma). The slides were then incubated in a blocking solution (PBS, 0.1% Tween-20, 3% BSA) for 30 min at RT.

### Microtubule shielding and immuno-staining

“Old” microtubules were revealed by blocking the access of anti-tubulin antibodies to the “young” microtubule containing the injected biotinylated tubulins. The shield consisted of four dense layers of antibodies. All incubation steps were performed for 30 min at room temperature. They were all separated by 3 washing steps in PBS supplemented with 0.1 of Tween-20. First, cells were incubated with anti-biotin antibodies (Jackson Immuno Research, ab2337630) at 1:50, followed by a secondary bovine antibody against goat IgG and labelled with Alexa Fluor 647 (Jackson Immuno Research, 805-605-180) at 1:50. When necessary, lattice shielding was preceded by PTM immunostainings. Detyrosinated and acetylated tubulins were labelled using anti-detyrosinated alpha tubulin (Abcam, AB48389) at 1:500 and anti-acetylated tubulin (Sigma, T7451) at 1:10000, followed by incubation with secondary donkey antibodies against rabbit IgG labelled with Alexa Fluor 488 (Jackson Immuno Research, 711-545-152) at 1:300 and donkey antibodies against mouse IgG labelled with Alexa 555 (Thermo Fisher, A31570) at 1:300. Four layers of antibodies were then added, by alternating layers of goat antibodies against bovine IgG (Jakson Immuno Research, 101-005-165) at 1:50, and bovine antibodies against goat IgG (Jackson Immuno Research, 805-005-180) at 1:50. Finally, the “old” MTs were revealed by immuno-staining them with monoclonal mouse antibodies against alpha tubulin (Sigma, T9026, DM1A) at 1:300 followed by a last incubation with goat antibodies against mouse IgG labelled with Alexa Fluor 405 (Thermo Fisher, A31553). Finally, the dishes were rinsed twice in PBS supplemented with 0.1% Tween-20, once in PBS, once in milliQ water and mounted in Mowiol 4-88.

### Imaging

Most immunofluorescence images were acquired using a confocal spinning disk microscope (Nikon Ti Eclipse) equipped with a spinning scanning unit (CSU-X1 Yokogawa), an ILAS 2 (GATACA) laser illumination system, and a R3 retiga camera (QImaging). Images were acquired using a Nikon Plan Apo l 60x/1.40 NA oil objective. Each wavelength was acquired separately with a 400nm Z-step width. Metamorph software was used for images acquisition.

Immunofluorescence images of figure 5 were acquired using an epifluorescence microscope (Nikon Ti Eclipse) equipped with a Prime BSI Express camera (Teledyne) and a Xcite LED1 (Excellitas) illumination system. Images were acquired using a Nikon Plan Apo VC 60x/1.40 NA oil objective. Micromanager 2.0 software was used for images acquisition.

### Quantification & analysis

Aquired images were analyzed using custom macros in Fiji (Schindelin et al., 2012). To determine microtubule ages in Figure 1 and in Figure 4, z-stack of each channel containing old (non-shielded) tubulin signal and new (injected) tubulin signal were first max projected. On each channel, an intensity-based threshold was manually applied to segment microtubules. For each channel, the average intensity of three single microtubule segments manually defined were used to normalized the raw image of each channel, then the corresponding segmented regions were used to measure the total normalized intensity on a summed projected image of the original z-stack. The ratio injected tubulin / total tubulin was computed as the total normalized intensity of the injected tubulin divided by the sum of the total normalized intensity of both injected tubulin and non-shielded tubulin.

### Image normalization and microtubule segmentation

To assess microtubule age or microtubule modifications (with or without nocodazole treatment), individual cells were cropped from the field of view, and each channel of the raw image was normalized independently. For each pixel, normalized intensity was calculated as I_norm = (I_px − I_bg) / (I_obj − I_bg), where I_obj was the average intensity of four lines drawn manually along individual microtubules (or modified segments), and I_bg was the average intensity of two regions drawn outside the cell to estimate background.

Microtubule segmentation was performed using Labkit classifiers trained on manually annotated images. For the microtubule-age experiments, a single classifier was trained on a stitched image combining the non-shielded and injected tubulin channels from one representative cell, and then applied to both channels in all cells. For nocodazole-treated cells, a single classifier was trained on a stitched image of the tubulin channel from five representative cells (one per treatment time point) and applied to all cells. Classifiers for acetylation and detyrosination were trained analogously, using stitched images of the corresponding modification channel from a small number of representative cells. In all cases, both individual objects and bundles of objects (microtubules or modifications) were annotated as foreground, while regions outside the cell and within the cytoplasm were annotated as background. Annotation and training were performed on full z-stacks to leverage axial information and improve segmentation accuracy.

All channels were segmented with their respective classifiers using a custom Fiji macro. Resulting probability maps were thresholded at 0.5 and 0.9 to generate object masks; masks obtained at the 0.9 threshold were used for subsequent analysis. For microtubule age quantification, the overlap between injected and non-shielded tubulin masks was subtracted from each, yielding masks referred to as “new-” and “old-”, respectively, each representing microtubules segmented exclusively in one channel. The overlap between these masks and the modification mask was then computed to identify modified microtubule segments.

### Intensity and length quantification

Normalized intensities were measured on the resulting masks slice-by-slice and summed across the z-stack to obtain total normalized intensity per cell for each network category: new-, modified-and-new-, old-, and modified-and-old-. For each category, the ratio of modified to total network intensity was calculated to estimate the proportion of modified microtubules.

Network length was estimated by converting total normalized intensity into micrometers using an empirical conversion factor. This factor was derived from some representative single-microtubule lines used during normalization. Line length was calculated from camera pixel size and imaging magnification. For each line, line intensity was measured on the segmented normalized image within a volume obtained by dilating the line and extending three slices above and below its z-position, to account for out-of-focus segmentation. Dividing this intensity by line length yielded a lineic intensity value; the empirical conversion factor was obtained by averaging lineic intensity values across all measured lines.

## Data availability

The data are available from the corresponding author upon request.

## Acknowledgments

This work was supported by fundings from the Agence Nationale pour la Recherche (ANR grant AAPG2022-PRC-SHARP and ANR-23-CHBS-0013) and the European Research Council (ERC consolidator grant 771599). Tanguy Chocat received a PhD fellowship from the Commissariat à l’Energie Atomique et aux Energies Alternatives (CEA). This work benefited from the technical contribution of the joint service unit CNRS UAR 3750. The authors would like to thank the engineers of this unit for their advice during the development of the experiments.

## Author contributions

(according to CRediT’s 14 Contributor Roles):

Tanguy Chocat: investigation, data curation, formal analysis.

Benoit Vianay: data curation, formal analysis, software, visualization.

Flora Silberzan: investigation.

Jérémie Gaillard: methodology, resources.

Louise Bonnemay: methodology, visualization.

Alexandre Schaeffer: methodology, supervision, validation.

Laurent Blanchoin: funding acquisition, supervision.

Manuel Théry: conceptualization, funding acquisition, investigation, supervision, validation, visualization, writing original draft.

## Disclosures

Authors declare that they have no competing interests.

---

**Figure S1.**
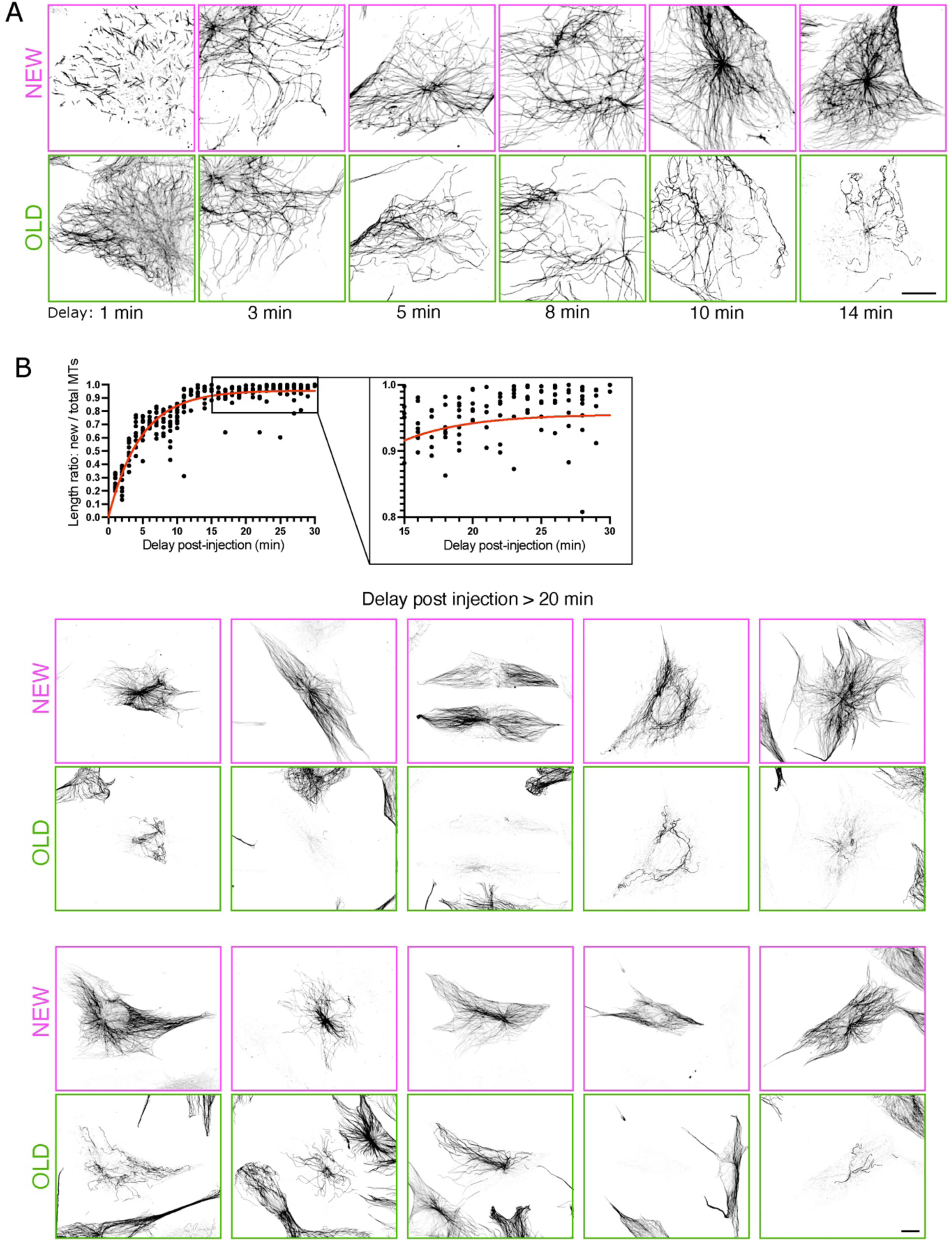
Detailed turnover of the MT network. (A) Images show examples of « new » MTs, assembled during the delay after tubulin injection, and « old » MTs, pre-existing to the injection and maintained during the delay, for 1, 3, 5, 8, 10 and 14 minutes long delays in cycling MEF. In each condition, « new » MTs are younger than the delay and « old » MTs are older than the delay. Scale bar represents 10 µm. (B) The graph shows a zoom of the graph shown in Figure 1E, highlighting the relative proportion of « new » MTs, and thereby the complementary non renewed and « old » MTs, 20 minutes after tubulin injection. Images show examples of such cells, in which « old » MTs are older than 20 minutes. Scale bar represents 10 µm.

**Figure S2.**
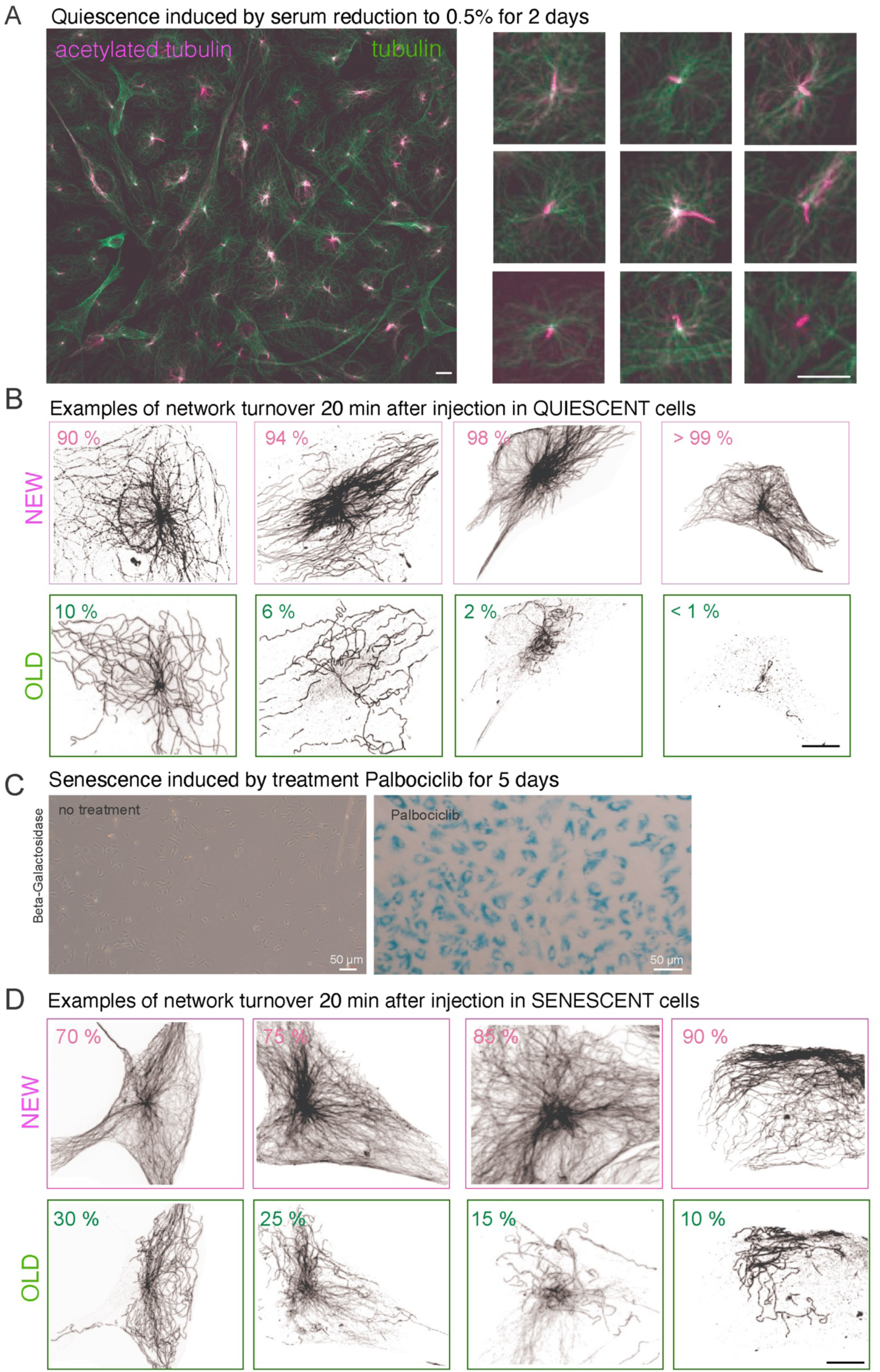
Detailed of the slowly-renewing MTs in quiescent and senescent cells. (A) Primary cilia on starved MEF cells. Left image shows a large-scale image of cells stained for acetylated tubulin (magenta) in order to detect primary cilia among the rest of the MT network (green). Rights images are zooms on individual cells to show the systematic presence of cilia. Scale bar corresponds to 10 µm. (B) Images show examples of injected quiescent cells, in which « old » MTs are older than 20 minutes. Scale bar represents 10 µm. (C) ß-galactosidase staining on cells treated or not with 12 µM Palbociclib for five days. Blue staining on the right image is characteristic of a positive staining of senescent cells. Scale bars represent 50 µm. (D) Images show examples of injected senescent cells, in which « old » MTs are older than 20 minutes. Scale bar represents 10 µm.

**Figure S3.**
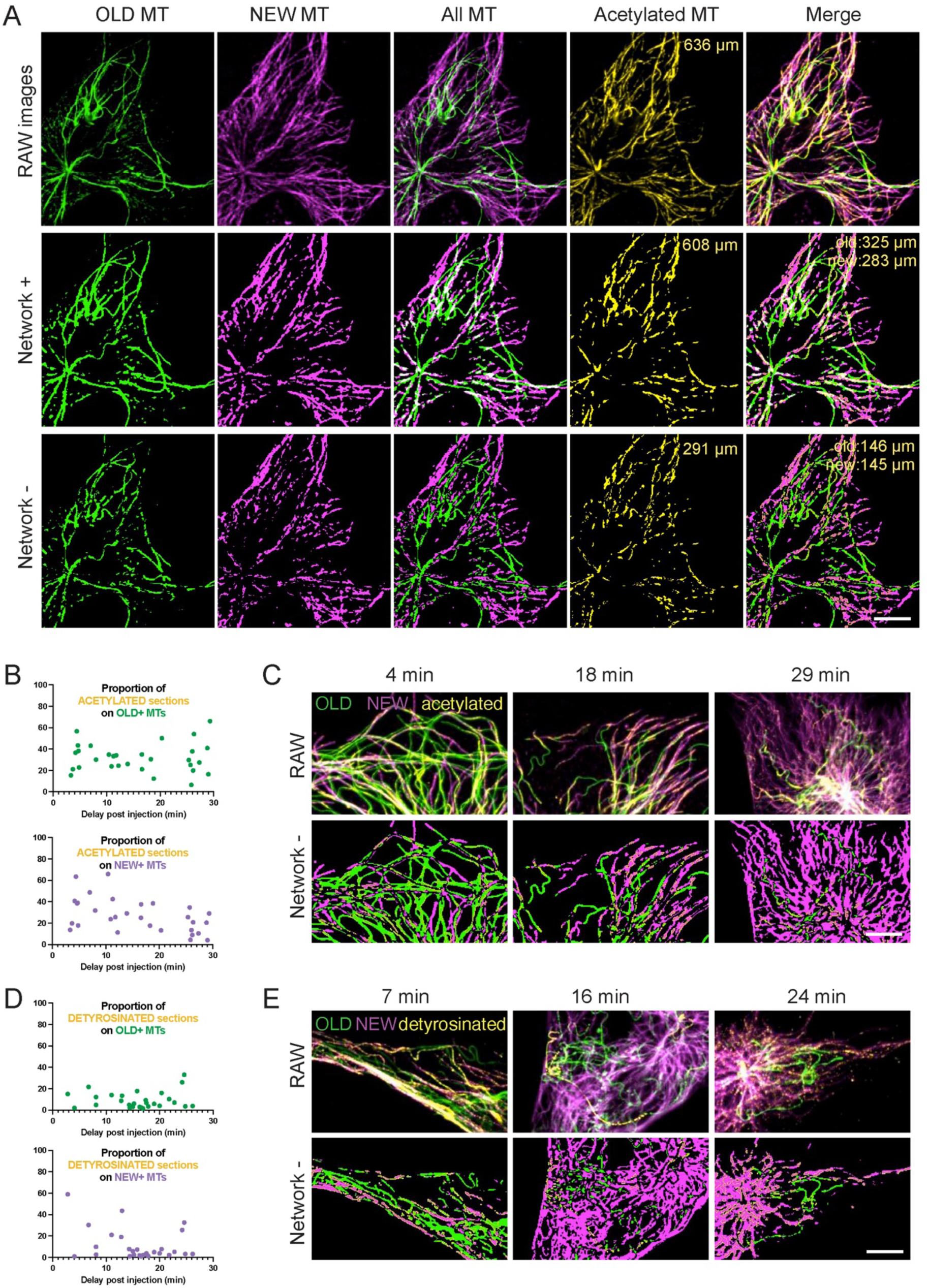
Network segmentation and quantification of PTMs. (A) The segmentation of the old (green) and new networks (magenta) using labkit led to two masks that were partially overlapping since some old and new MTs occupied the same pixels. When overlaying the entire networks, named “network+”, we could visualize the overlap regions (in white in the central column). We could also substract these common pixels in order to generate smaller networks, named “network-”, which showed no overlap (see the absence of white in the central image of the third line). Combining these “network+” or “network-” with the segmentation of the acetylated sections (yellow contours), we could calculate the upper and lower boundaries of the length of the modified network. The “network+” led to the high boundary since some acetylated pixels could actually belong to the other network. The “network-” led to the low boundary since the pixels belong exclusively to this network. Values in yellow indicate the total length of the acetylated sections in the corresponding networks. Scale bar represents 10 µm. (B) Graphs show the length of the « new » and « old » networks that were acetylated relative to the entire « new » and « old » network, depending on the delay after injection, ie the age limit between these two populations. In these graphs the lengths were estimated on the the “network+”, ie the entire network, so values were upper-estimation of the actual modified length. Underestimated values, calculated from the “network-”, ie the old and new networks without any overlap, were shown in figure 3B. (C) Images show examples of segmentation of acetylated regions (yellow contours) overlapping with old and new MTs for various delay after injection. Acetylated sections were visible on 4-minute-old MTs and some 29-minute-old MTs were not fully acetylated. Scale bar represents 10 µm. (D) Graphs show the length of the « new » and « old » networks that were detyrosinated, depending on the delay after injection, ie the age limit between these two populations. In these graphs the lengths were estimated on the the “network+”, ie the entire network, so values were upper-estimation of the actual modified length. Underestimated values, calculated from the “network-”, ie the old and new networks without any overlap, were shown in figure 3F. (E) Images show examples of segmentation of detyrosinated regions (yellow contours) overlapping with old and new MTs for various delay after injection. Detyrosinated sections were visible on 7-minute-old MTs and some 24-minute-old MTs were not fully detyrosinated. Scale bar represents 10 µm.

## Notes

### Competing Interest Statement

The authors have declared no competing interest.

